# Harnessing NCX-IP_3_R-dependent Calcium Oscillations to Regulate Angiogenic Signaling in Endothelial Cells

**DOI:** 10.64898/2026.04.07.717042

**Authors:** Donghyun Paul Jeong, Stephen Cini, Kailee Mendiola, Satyajyoti Senapati, Alexander Dowling, Hsueh-Chia Chang, Jeremiah Zartman, Donny Hanjaya-Putra

**Affiliations:** Bioengineering Graduate Program, University of Notre Dame, Notre Dame, IN 46556; Aerospace and Mechanical Engineering, University of Notre Dame, Notre Dame, IN 46556; Chemical and Biomolecular Engineering, University of Notre Dame, Notre Dame, IN 46556; Harper Cancer Research Institute, University of Notre Dame, Notre Dame, IN 46556; Center for Stem Cell and Regenerative Medicine, University of Notre Dame, Notre Dame, IN 46556

**Author notes:** **Address correspondence to:** Donny Hanjaya-Putra, 2010G McCourtney Hall East, University of Notre Dame, Notre Dame, IN 46556.

## Abstract

The blood vasculature has a high capacity for structural regeneration, driven by the blood endothelial cells (BECs) that comprise it. This regenerative process, which involves BEC migration and proliferation to form these complex tissues, is linked to low frequency (< 0.1 Hz) calcium spiking that precedes these activities. However, we need new approaches to stimulating angiogenic responses in tissue engineering applications. By conducting experiments that manipulate local ionic concentrations and developing a simple, yet powerful, computational analysis, we demonstrate that sodium-calcium cross-talk is a crucial component that regulates the calcium signaling and downstream angiogenic responses. Activation and deactivation of the inositol triphosphate 3 receptors (IP_3_Rs) on the endoplasmic reticulum (ER) and the switch between forward and reverse modes of the sodium-calcium exchanger (NCX) are proposed to be the key mechanisms underlying calcium oscillations when cells are exposed to temporary cationic depletion. The spiking is suggested to be a release of intracellular calcium mediated by IP_3_R activity, and transport in or out of the cell is driven by NCX for the calcium oscillatory signaling pattern. The NCX and IP_3_R both contribute to manage intracellular calcium and ionic concentration as initially there is a long ER deactivation period while intracellular sodium slowly increases until a sudden onset of calcium is released by the ER. Other calcium and sodium ion channels can change this resonant coupling of ER and NCX to alter the inter-spike duration. Synchronization of the spiking intervals between cells is triggered by stimulating with vascular endothelial growth factor (VEGF), which induces a propagating wave of intracellular calcium across the 2D tissue culture, prior to coordinated cell migration and proliferation towards the VEGF source. This wave, which can be artificially induced and studied using electrical stimulation, suggests that the underlying sodium-calcium crosstalk mechanism introduces intracellular calcium polarization, whose orientation is transferred across cells through spike synchronization. Thus, control of calcium signaling dynamics through regulation of ionic depletion can serve as useful method for generating angiogenic responses in engineered tissue constructs.

## 1. Introduction

Intracellular calcium signaling plays a crucial role in many key cellular processes, such as proliferation or apoptosis, metabolism, secretion and gene regulation, and cell motility.^1–4^ In response to external signals, cells are capable of drastically increasing the flux of calcium ions from either extracellular space or intracellular calcium storage in organelles such as endoplasmic reticulum (ER).^5,6^ Calcium signaling is a ubiquitous aspect of many intracellular communication pathways because of a large set of signals, receptors and transporters, together known as “calcium signaling toolkit”, that can be applied to multiple cellular processes.^7,8^

In some cells, calcium-based signaling works at time scales in the order of milliseconds to seconds, allowing rapid fluctuations in calcium signaling. Excitable cells, such as cardiomyocytes, neurons, and muscle cells, can have frequency of between 1-100 Hz.^9,10^ Calcium also plays a role in slower dynamics that result from a gradual buildup of calcium over time, which can influence gene expression.^11^ In between these two cell types are the types of cells that undergo sharp calcium oscillations similar to excitable cells, but at a significantly slower timescale. These are electrically non-excitable cells that undergo oscillation in the frequency range of between 0.1 and 0.001 Hz.^9^

Because calcium activity is often transitory and occurs across a broad range of frequency domains, imaging-based calcium measurement are most commonly used to map out calcium dynamics. Calcium indicators, such as Fura-2 dye, results in excitation shift or fluorescence intensity when bound to calcium.^12^ Alternatively, reporter cell lines expressing a genetic encoded calcium indicator exhibit a fluorescence shift with calcium concentration fluctuations. Overall, calcium measurements rely on fluorescent imaging-based techniques, as it is best suited for live cell imaging at high sampling rates. Understanding calcium signaling is especially important in tissue engineering and other bioengineering applications that focus on population-wide cell communication. Calcium, in addition to playing an important role in intracellular signaling, also serves as an intercellular communication tool in many cells, aiding in spontaneous self-organizational behavior of cells often seen in organogenesis.^8,13–15^

In constructing engineered tissues, spontaneous self-organization is often the only way to form structures such as microvasculature that are smaller than the minimum spatial resolution of devices such as 3D bioprinters.^16^ To induce self-organization, heterogeneous cell population can be seeded onto a matrix and stimulated with relevant growth factors.^17,18^ Cells can respond to mechanical stimuli such as shear stress, chemical gradients of nutrients and growth factors, or contextual cues such as presence of other cells near it.^19–21^ In fact, numerous studies have indicated that the spatial distribution and intercellular communication play crucial roles in the self-organization of the cells into functional tissues.^22,23^

This calcium signaling toolkit is especially relevant in vascular tissue engineering. Calcium spiking plays a crucial role in directing endothelial cellular response to various external stimuli, such as growth factors, mechanical forces, or metabolic stresses, that that are present in a vascular system.^24^ For example, shear stress is known to induce cytoplasmic calcium spiking with a response time in the range of 1 minute in blood and lymphatic endothelial cells.^25–27^ Endothelial cells also exhibit repeated calcium oscillation in response to reoxygenation following a period of hypoxic stress.^28^ Most relevant to the tissue engineering application, however, is the calcium-mediated endothelial response to pro-angiogenic growth factors, most famously vascular endothelial growth factor (VEGF). VEGF is known to induce physiologically relevant responses in endothelial cells, such as proliferation and migration, that are crucial for angiogenesis.^29,30^ Previous studies have demonstrated that VEGF induces calcium signaling dynamics via amplitude modulation or frequency modulation.^31^ However, due to the lack of high-throughput and robust calcium data quantification methods, combined with the high degree of heterogeneity in spiking behavior among endothelial cells, it remains difficult to obtain quantitative measurements of subtle frequency shifts in response to external stimuli.

Therefore, it is clear that there is a need for a better understanding of how calcium dynamics in large non-excitable cell populations affect angiogenesis. Using newly developed computational analysis tools, we identify a previously uncharacterized calcium-signaling pathway that modulates endothelial responses to angiogenic growth factors. We quantify key metrics describing endothelial responses to VEGF and reproduce these effects through controlled ionic depletion using a novel microfluidic device. Finally, we introduce a mathematical model based on sodium-calcium exchanger and IP3R-mediated calcium fluxes that captures endothelial calcium oscillations and predicts cellular responses to external perturbations. Altogether, our study suggests that control of calcium signaling dynamics through regulation of ionic depletion can serve as useful method for generating angiogenic responses in engineered tissue constructs.

## 2. Results

### 2.1 Automated computational pipeline for large-scale calcium oscillation data extraction in non-excitable cells

We designed a data processing pipeline, as shown in **Figure 1**, that is designed around analyzing the intracellular calcium oscillation patterns in non-excitable cells. The overall pipeline receives the raw fluorescent image stack at a magnification between 4 & 10x in tiff format. These images capture the fluorescent signal intensities of the mesodermal-lineage cells which corresponds to cytoplasmic calcium concentration. For this study, we relied on Calbryte 520AM dye, a robust calcium indicator dye that is non-fluorescent until bound with calcium ions within the cell.

**Figure 1.**
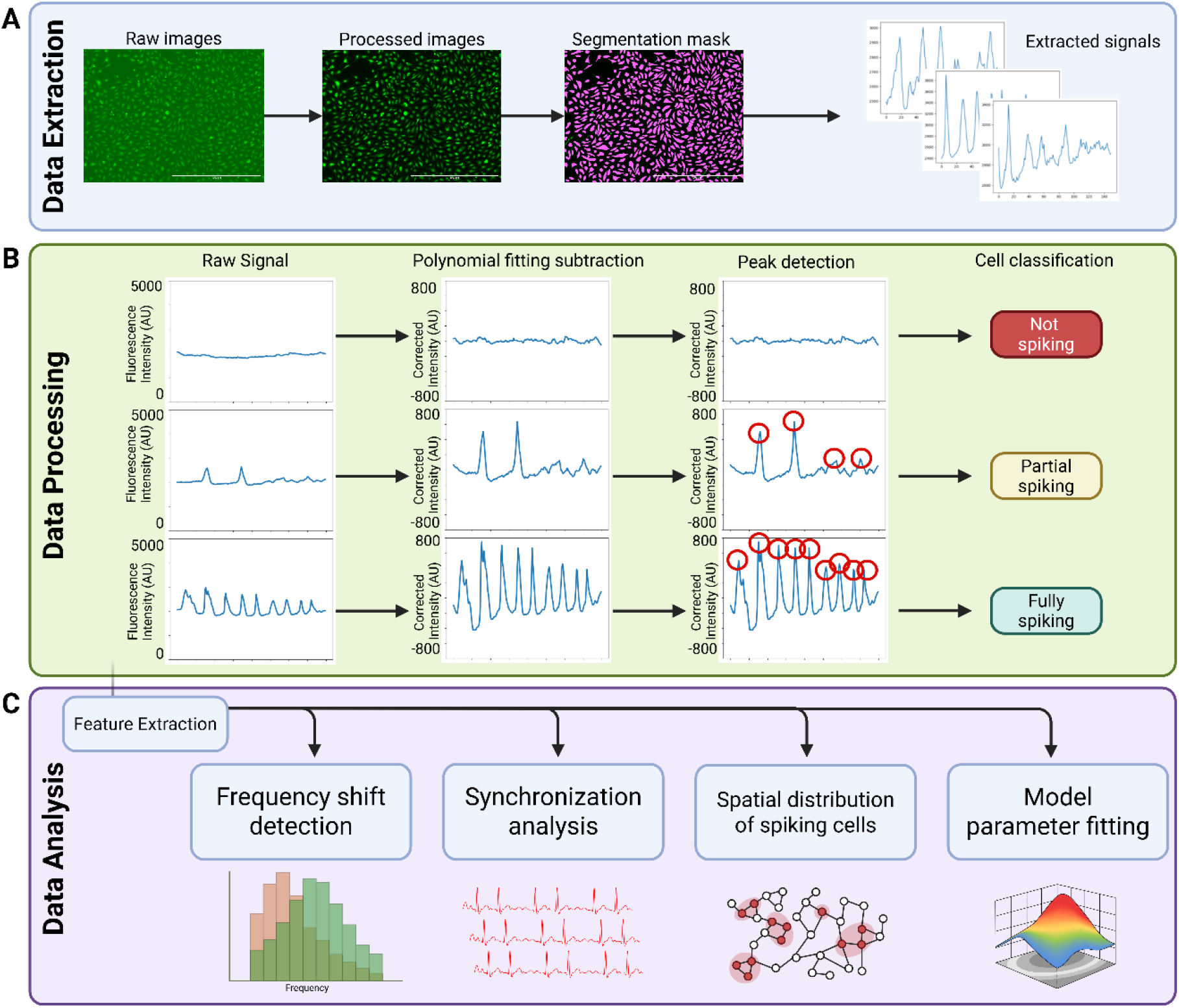
Three unique subtypes of endothelial calcium oscillation patterns captured by the automated calcium analysis. (**A)**. Steps for initial image data processing and extraction. **(B)**. Extracted temporal calcium data is further processed and filtered. **(C)**. The processed oscillatory data is used for various analysis downstream, such as frequency shift or synchronization. Scale bars represent 1 mm.

Before proceeding with the pipeline, the images require undergoing a preprocessing step **(Figure 1A)**. This process begins with a rolling ball background subtraction to remove the effects of uneven background signal. It is followed by a despeckle step to remove the signals attributed to non-cellular fragments. Next, a dead cell detection step follows, identifying and excluding auto fluorescent dead cells by setting a maximum fluorescence intensity threshold. The processed images are then segmented using either a Cellpose-based algorithm or a traditional particle detection algorithm. This segmentation is performed on the max-projection image generated from the entire image stack, with optional image drift correction applied as needed. The resulting segmentation masks are used to generate the average intensities for each cell at every timepoint, which is saved to an intensity (n-by-t) matrix, where n in the number of identified cells and t is the total number of timepoints.

After preprocessing the image stack, the next step is data processing **(Figure 1B)**. The extracted data is then processed individually by fitting a 6^th^ order polynomial to each timeseries data, then subtracting the values by the fitted polynomial to correct for the overall background signal increase or loss of fluorescence due to photobleaching. Afterwards, the cells are classified into either spiking, partial spiking, or non-spiking cells based on a peak detection algorithm, which identifies the timepoints at which the signature calcium spike peaks occur. The parameters of the peak thresholding are set by the user based on the imaging setup and the cell behavior. The pipeline takes the processed data for spiking cells and extracts parameters for subsequent analysis such as frequency shift or synchronization (**Figure 1C**). These data can be exported and analyzed for correlation with the cell morphological or spatial information identified from the corresponding cell masks extracted from the cell segmentation step.

We demonstrate the viability of our method in detecting the irregular calcium oscillation patterns in non-excitable cells. The sample images consist of human dermal blood endothelial cells (BECs) which were cultured to confluency. They were stained with Calbryte 520AM dye without probenecid treatment to capture the calcium dynamics of cells. A resulting image at a single timepoint is shown with representative images of each cell type (spiking, partial spiking, or non-spiking) identified by the pipeline **(Supplementary Figure S1A)**.

We compare the results of our algorithm with the manual segmentation method, which consists of manually segmenting and identifying the spiking cells. We demonstrate that the automated method and the manual segmentation can identify approximately similar number of cells, but with greater consistency with automation **(Supplementary Figure S1B-C)**. The calcium data for the two extraction methods both estimated similar oscillating period and percentage of spiking cells, demonstrating that there is no significant difference between the two methods **(Supplementary Figure S1D-E)**. Taken together, these results demonstrate that our automated algorithm can reliably capture highly irregular calcium spiking dynamics in a large cell population in a timely and non-labor-intensive manner.

### 2.2 Endothelial cells exhibit a robust acute and chronic calcium response to VEGF

By utilizing the pipeline approach to characterize the calcium behavior within non-excitable cells, we reliably capture the calcium frequency shifts seen with the addition of VEGF. First, we observed that the addition of VEGF causes an immediate wave of sudden spiking of calcium concentration that propagates through the endothelial population within order of seconds following the addition **(Figure 2A, Supplemental Movie S1)**. We estimate this wavefront of calcium spike activity propagates at an approximate rate of 2400 μm/min, calculated using the timepoints at which cells across the imaging window reached maximum calcium intensity **(Figure 2B)**. This rapid propagation indicates that the wavefront is independent from diffusion of VEGF molecules, which has an expected diffusive velocity in the order of μm/min, assuming a diffusivity of 10^-6^ cm^2^/s over a few cm of the imaging window. Therefore, we hypothesized that the VEGF-mediated wave propagation is most likely due to intercellular exchange of ions (e.g., through gap junctions) rather than VEGF diffusion.

**Figure 2.**
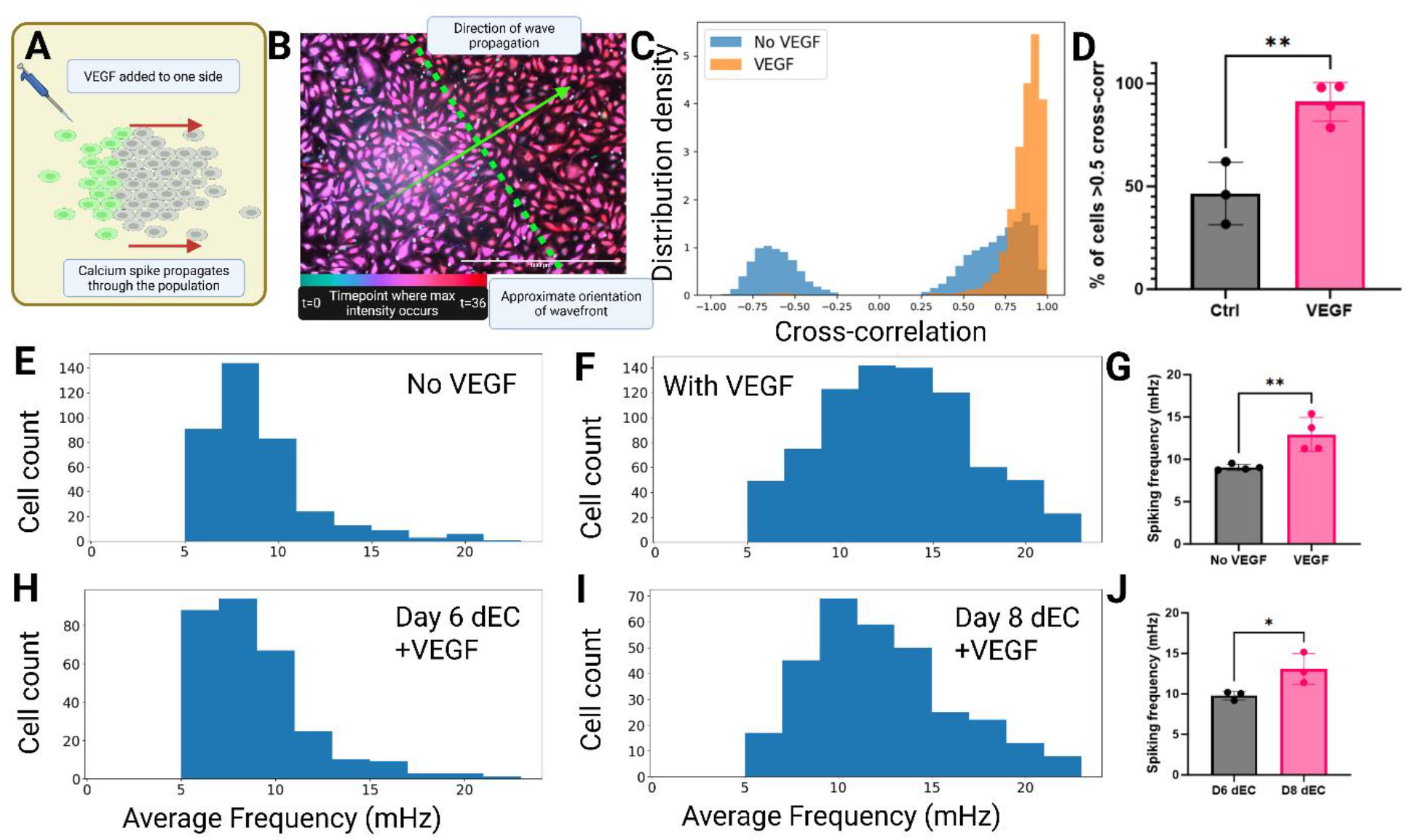
VEGF induces a frequency shift in calcium-oscillatory endothelial cell population. **(A)** Schematic of the VEGF-induced calcium wave propagation within seconds of adding VEGF. **(B)**. Temporal color code image of the cells following the VEGF addition. The color corresponds to the timepoint at which the maximum intensity is reached. An approximate wavefront is shown in green dotted line, and the direction of wave propagation, which is perpendicular to the wavefront, is shown in green solid line. **(C)**. Distribution density plot of the degree of cross-correlation for each cell-cell pair within the imaging window for VEGF and no-VEGF conditions. **(D)**. Percentage of cell-cell pairings that exhibit higher than 0.5 in cross-correlation value, n=3 for control and n=4 for VEGF. **(E-F)**. Histogram showing the distribution of average oscillation period in oscillatory cells cultured with and without VEGF. The cells with VEGF were subjected to 50ng/mL of VEGF for 24 hours. **(G)**. Average spiking frequency of cells treated with and without VEGF, n=4. Each run was with the identical donor but cultured and measured independently. **H-I**. Average oscillation period histogram for iPSC-differentiated ECs at day 6 and day 8 of differentiation. Mean oscillation periods are shown on the top right for each plot. **J**. Average spiking frequency of iPSC-dEC at day 6 and 8, n=3 for both groups. *p<0.05, **p<0.01. Scale bar represents 1mm.

We first quantified the degree of synchronization by measuring the cross-correlation values for each cell-cell pair within the imaging window. Cross-correlation values were calculated assuming no temporal shifts. When the VEGF-stimulated cells were compared against the non-stimulated cells, we observed a clear shift towards higher synchronization, with the number of cell-cell pairs exhibiting at least 0.5 cross-correlation value from around 50% to nearly 100% **(Figure 2C-D, Supplementary Figure S4)**.

We then collected calcium measurement 24 hours after exposure to VEGF and measured the average spiking frequency for each cell exhibiting oscillatory behavior. In a non-VEGF treated condition of less than 0.5 ng/mL of VEGF, the cells exhibited approximately 8 mHz spiking frequency **(Figure 2D, F, Supplementary Movie S2)**. This spiking frequency shifted to approximately 12 mHz with the addition of 50 ng/mL of VEGF **(Figure 2E, F, Supplementary Figure S2, Supplementary Movie S3)**. This pattern was conserved with treatment as low as 10ng/mL of VEGF with no dose-dependent changes to frequency, suggesting that VEGF-activated frequency shift is a highly sensitive stepwise activation mechanism **(Supplementary Figure S3)**.

Interestingly, we also found that induced pluripotent stem cell (iPSC)-derived endothelial cells exhibit this same frequency as they reach terminal differentiation. Human iPSCs are typically differentiated to endothelial cells through a stepwise treatment involving various growth factors, followed by maturation into mature endothelial cells through VEGF treatment. We performed differentiation of iPSC to endothelial cells using an 8-day stepwise protocol and measured the calcium spiking frequencies at day 6 and day 8. As expected, at day 6, the immature differentiated endothelial cells (dEC) exhibited a low spiking frequency of around 8 mHz even in the presence of 50 ng/mL of VEGF **(Figure 2G, I, Supplementary Movie S4)**. In contrast, fully matured dECs at day 8 exhibited a 11 mHz spiking frequency, which matched the response of the native blood endothelial cells in the presence of VEGF **(Figure 2H, I, Supplementary Movie S5)**. These results suggest that spiking frequency response defines the degree of maturity of iPSC-derived ECs.

### 2.3 Electrical stimulation directly induces acute calcium responses in ECs

To show that we can mimic the effects of VEGF through regulation of external ionic concentration, we devised a microfluidic channel equipped with a selectively permeable ionic membrane (SPIM) which was previously published.^32^ We modified a single channel fluidic chip with two wells by inserting a narrow pipette tip within one of the wells **(Figure 3A)**. The pipette tip was modified by inserting the SPIM to the narrow opening such that the membrane fully separates the inner-pipette chamber and the outer well of the chip. Then, a negative electrode was placed inside the media in the pipette tip, and a positive electrode was placed in the outer well. When the current was applied, we confirmed that the charged dye placed inside the microfluidic channel is transferred to the pipette tip through the SPIM **(Figure 3B)**. Using this microfluidic chip, we synchronously depleted the cations present in the cell culture media, which noticeably shifted in the degree of synchronization in BEC oscillation similar to that of VEGF stimulation **(Figure 3C, Supplementary Movie S6)**.

**Figure 3.**
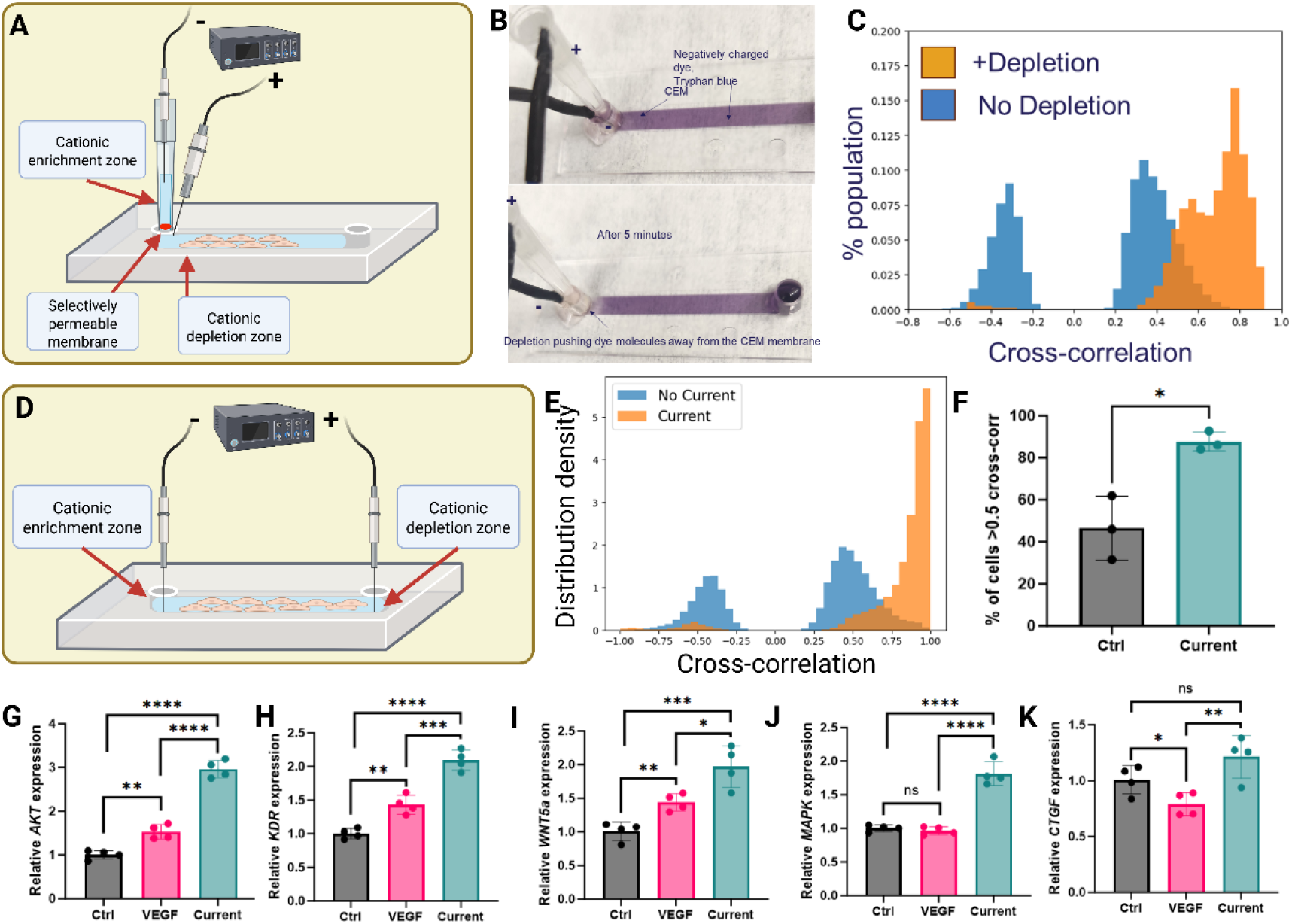
VEGF or electrical stimulation induces intercellular calcium waves across coupled endothelial cells. **A**. Schematic for the cation depletion setup, showing the placement of a double-well with a selectively permeable membrane. **B**. Demonstration of the viability of the depletion chip showing the removal of negatively charged Tryphan Blue dye after 5 minutes of applying the current. **C**. Histogram showing the distribution of degree of cross-correlation for each cell-cell pair before and after cationic depletion. **D**. Schematic of the current-induced transient ionic depletion in a microfluidic channel. **E**. Distribution density plot of the degree of cross-correlation with and without the current applied. **F**. Percentage of cell-cell pairings that exhibit higher than 0.5 in cross-correlation value, n=3 for both groups. **G-K**. Relative RNA expression level changes for *AKT, KDR, WNT5a, MAPK*, and *CTGF* with VEGF treatment and current treatment normalized to the non-treated cells, run in technical replicates of n=4. *p<0.05, **p<0.01, ***p<0.001, ****p<0.0001.

We then hypothesized that a temporary shift in external ionic concentration would mimic the transient effect of VEGF on cellular calcium signaling synchronization. Therefore, we devised a new microfluidic chip where two electrodes are placed in the two opposing wells, and a controlled current is applied directly through the cell culture area to induce ionic movement and a localized, transient cationic depletion zone **(Figure 3D)**. We optimized the current and voltage parameters required to mimic the effect of VEGF on calcium oscillation synchronization and identified a range of 10-12V and 4.5-5.5mA as optimal. Using these parameters, we applied a 50 second pulse of current to the BECs and measured the calcium oscillation pattern during and after the pulse, where the BECs exhibited the same synchronization behavior as seen with VEGF-stimulation **(Figure 3E, Supplementary Movie S7)**. Nearly 90% of cells treated with the current exhibited a cross-correlation coefficient of >0.5, which is significantly higher than their non-treated counterpart **(Figure 3F, Supplementary Figure S4)**.

We also explored whether this current-based calcium synchronization is sufficient to replicate the physiological effects of VEGF in cells. We compared the transcriptional levels of various VEGF-related genes between VEGF-treated, electric current-treated for a single 50 second pulse, and non-treated cells after 6 hours of initial stimulation. We found that the VEGF-treated cells had elevated levels of *AKT, WNT5A*, and *KDR* compared to non-VEGF treated cells, while the current-treated cells exhibited a significantly greater upregulation of the same genes at the 6-hour timepoint **(Figure 3G-I)**. *AKT* is a component of the *PI3K/AKT* pathway, regulating cellular processes such as proliferation and cell survival, and is a known target of VEGF-mediated signaling.^33–37^ *WNT5A* is endogenously expressed in endothelial cells and is involved in pro-angiogenic pathways.^38,39^ *KDR*, or VEGFR2, is the primary receptor of VEGF in endothelial cells and is crucial for VEGF-mediated signaling cascade.^40,41^ The parallel upregulation of these genes in the current- and VEGF-treated cells indicates that the current-stimulation can successfully recapitulates key downstream signaling effects of VEGF.

Notably, *MAPK*, another crucial VEGF-dependent pathway, was not significantly elevated in VEGF-treated cells but was significantly elevated in current-treated cells, which may indicate a temporal delay in VEGF-*MAPK* activation pathway that the current-based calcium stimulation is able to bypass **(Figure 3J)**.^42,43^ Conversely, *CTGF*, which can inhibit VEGF-induced angiogenesis, was downregulated in VEGF-treated cells, but the current-treated cells were unaffected. This finding suggests that the calcium synchronization works in a pathway specific to the VEGF signaling cascade **(Figure 3K)**.^44^ Taken together, our data suggests that the transient electrical stimulation or ionic depletion through the use of microfluidic-inspired circuits can mimic the effects of VEGF-induced angiogenesis through modulation of intracellular calcium.

### 2.4 Calcium spiking couples with predicted sodium dynamics through NCX transporter

Classical mathematical models such as the FitzHugh-Nagumo model or the more recent Ten Tusscher model describe calcium spiking in excitable cells through rapid, coupled changes in membrane potential.^45,46^ In electrically non-excitable endothelial cells, however, the membrane potential does not vary on the same time scale, necessitating a different mechanism to explain the observed oscillatory behavior **(Supplementary Movie S8)**. Because the plasma membrane is more permeable to potassium than sodium due to K-leak channels, potassium contributes more strongly to the membrane potential.^47,48^ This suggests that in non-excitable endothelial cells, calcium dynamics may be coupled more closely with sodium than potassium. We therefore hypothesized that small sodium fluctuations, coupled to calcium via sodium-calcium transporters, can generate large calcium spikes without producing comparably large changes in membrane potential.

To test this hypothesis, we measured the calcium oscillation dynamics before and after a sudden twofold increase in external sodium concentration from 140 mM to 300 mM.^49^ As shown in the representative calcium concentration trace, nearly all cells stop the oscillations abruptly just seconds after the sodium spike **(Figure 4A, Supplementary Movie S9)**. This lack of oscillation can be drastically reversed within seconds by reducing the sodium concentration to 140 mM again by diluting the media with deionized water (**Supplementary Movie S10**). In comparison, the cells continue to oscillate when the potassium concentration is increased tenfold from 5 mM to 50 mM, indicating that the calcium oscillation is primarily coupled specifically with sodium (**Supplementary Movie S11**).

**Figure 4.**
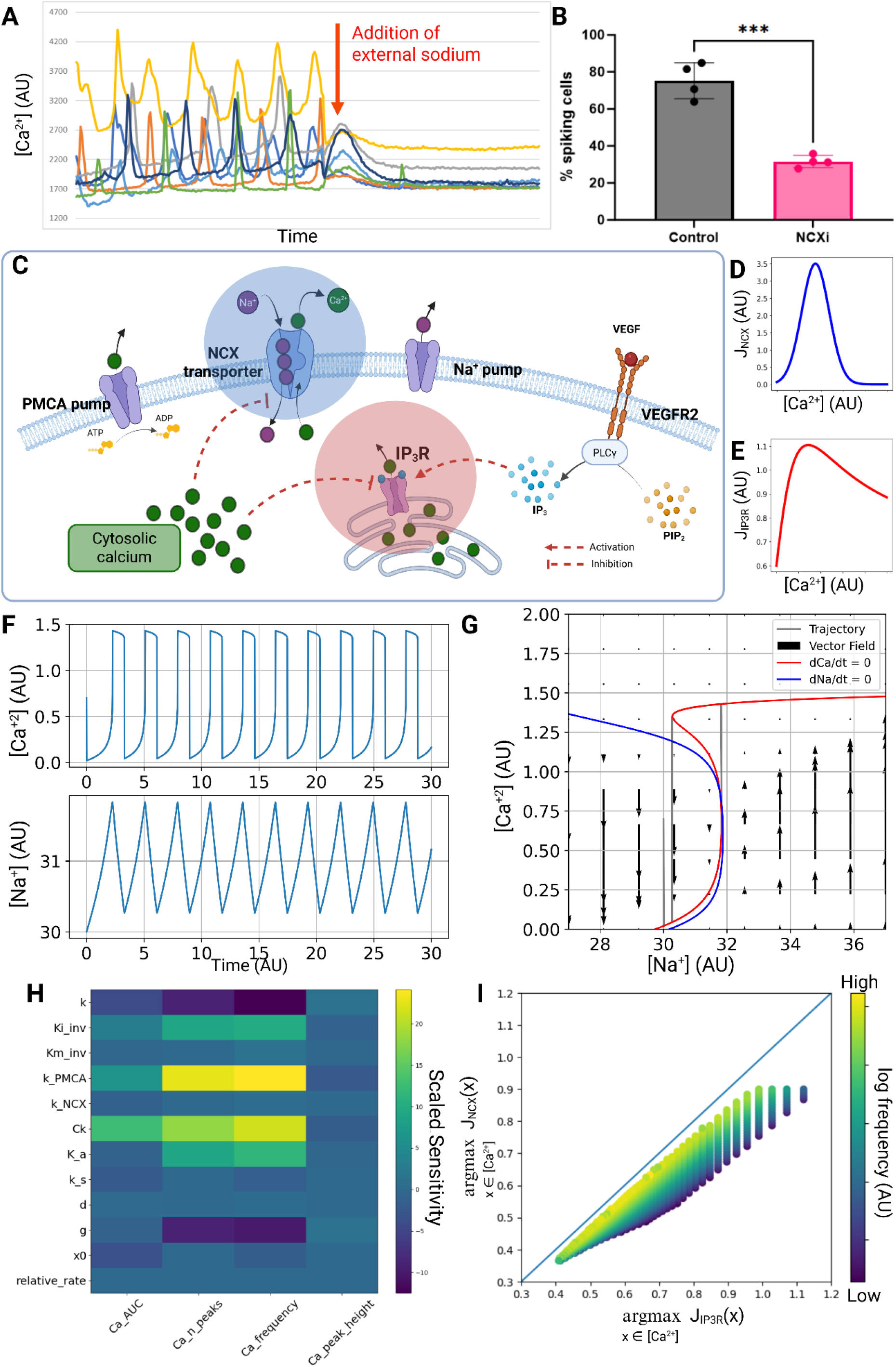
An IP_3_R-NCX calcium oscillation model can produce calcium oscillatory system in non-excitable cell model. **A**. Representative calcium intensity trace for one cell undergoing external sodium spike from 140mM to 300mM, which shows a sudden disappearance of all calcium oscillation activity, similar to calcium dynamics predicted in figure H. **B**. Percentage of oscillatory cells when treated with 10 μM SEA0400 NCX inhibitor showing significant inhibition of oscillatory behavior, n=4. **C**. Diagram showing the proposed mechanism of NCX-IP3R coupled calcium oscillatory system. The two key transporters with activation-inhibition curves with respect to calcium are highlighted with shaded circles. **D-E**. Calcium flux of NCX and IP3R transporters with respect to cytosolic calcium concentration showing maximum transport rates at similar calcium concentrations. **F-G**. Simulated calcium and sodium oscillation and the associated nullclines on Ca-Na phase plane showing a single unstable equilibrium in the middle branch of the S-hysteresis. Calcium nullcline is shown in red, sodium nullcline is shown in blue. **H**. Local feature sensitivity analysis of the model with respect to each parameter. Area under the curve (AUC), number of peaks, and frequency of the peaks for calcium are quantified for the sensitivity analysis. The values shown are for central difference analysis with ±1% perturbations. **I**. Effect on the frequency of the oscillation of the calcium concentration at the maximum flux for IP3R and NCX transporters. Color scheme corresponds to log of the frequency of the oscillation, while lack of points indicates a non-oscillatory system. Oscillation is only possible when the peak flux of IP3R is achieved at a slightly higher calcium concentration compared to that of peak NCX flux. ****p<0.0001.

We hypothesized that the sodium-calcium exchanger transporter (NCX) is the most likely coupling mechanism of sodium and calcium. NCX is a widely studied ion antiporter that transports one calcium ion for every 3 sodium ions.^50^ We treated the endothelial cells with 10μM of SEA0400, a known NCX-inhibitor, and found the percentage of cells undergoing oscillation decreased dramatically from around 75% to 25% **(Figure 4B, Supplementary Movie S12)**.^51^ Taken together, these results suggest that the calcium oscillations in non-excitable cells are coupled with sodium dynamics through NCX.

### 2.5 Mathematical model for calcium oscillation in non-excitable cells

Based on these insights, we developed a mathematical model to describe the calcium spiking patterns in non-excitable cells, which do not exhibit drastic changes to membrane potential compared to excitable cells. Our model relies on the NCX-IP_3_R-dependent calcium dynamics pathway **(Figure 4C)**.

It has been widely reported that the inositol 1,4,5-trisphosphate receptor (IP3R) is primarily responsible for the rapid release of calcium from the endoplasmic reticulum (ER).^52–54^ Importantly, IP3R exhibits a biphasic regulatory mechanism with respect to cytosolic calcium concentration (calcium-induced-calcium-release mechanism): its calcium release is activated by allosteric binding of calcium, but inhibited at saturating concentrations of cytosolic calcium.^55–57^ This widely studied mechanism of IP3R-mediated calcium release is critical for the oscillatory behavior of calcium. This biphasic behavior of calcium release can be modeled using Michaelis-Menten kinetics for substrate-inhibitory reaction as follows:

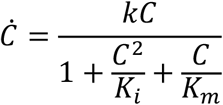

where *k* is the rapid ER calcium release kinetic constant and *K*_*m*_ and *K*_*i*_ are the association equilibrium constants of calcium binding for transport and substrate inhibition. The biphasic mechanism produces an optimum calcium concentration at 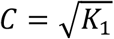 for maximum calcium ER release rate and the bandwidth of calcium concentration around this highest calcium release rate is governed by 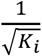.

Next, we model the calcium efflux through the NCX transporter. NCX transporter in typical physiological states is driven by the sodium gradient across the cell membrane, which allows the calcium to be depleted from the cytosol against its own concentration gradient.^58^ NCX also exhibits allosteric activation by intracellular calcium through its two binding domains, CBD1 and CBD2, which increases the calcium transport rate.^59–61^ However, very high levels of intracellular calcium can also lead to reversal of the NCX transport direction through autoinhibition or indirectly through calmodulin regulation or activation of NCX-cleaving protease.^62–64^ We model this behavior through a simple bell curve with the peak transport rate achieved at a set calcium level, and the transport rate driven by sodium gradient, using the following rate term:

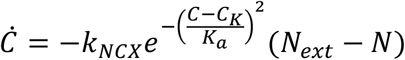

where C_K_ represents the critical concentration of calcium at which the maximum NCX transport rate is reached, and K_a_ represents the rate at which the calcium binding with the allosteric activation and inhibition sites are achieved. This calcium/sodium efflux/influx rate has a maximum at an intracellular calcium concentration of *C* = *C*_*k*_.

Both the key ER calcium release rate and the key NCX calcium/sodium efflux/influx rate have distinct maxima at particular intracellular concentrations. These two terms both exhibit optimal Ca concentrations. When these two optimum concentrations align, the calcium dynamics is expected to slow down to the level of the sodium dynamics, when the intracellular calcium concentration is away from the aligned values. When the calcium concentration reaches the optimum value, rapid calcium dynamics takes over and a universal spiking duration is observed. Other ion channels serve to shift this alignment and hence the interval between spikes. We will use a null-cline analysis to explore the dynamics when the optimum concentrations for peak ER/NCX rates are nearly aligned **(Figure 4D-E)**.

We further add a simple transport rate constant for plasma membrane calcium pump (PMCA) which is capable of pumping calcium at a low but steady rate even at low calcium concentrations driven by ATP.^65^ Therefore, we propose the following two for the dynamics of the intracellular calcium concentration C and the intracellular sodium concentration N:

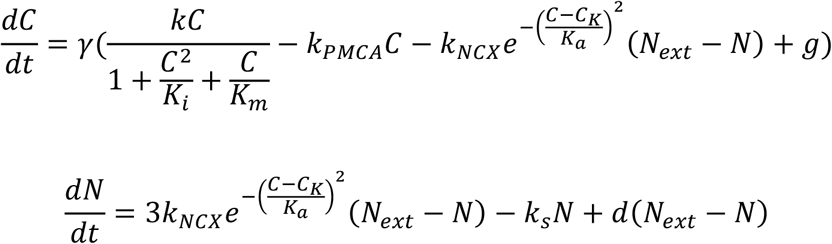

where *k*_*s*_*N* term represents the constant sodium pumping achieved by ubiquitous sodium pumps, d represents the concentration gradient-driven rate of sodium leakage across the plasma membrane, and *g* represents a constant calcium leakage from the ER.^66^

At equilibrium, the intracellular sodium concentration N is 3 orders of magnitude higher than that of the calcium C. Hence, the sodium flux terms must nearly cancel each other in the dynamics for the sodium ion to produce slow dynamics. Consequently, the related kinetic constants *k*_*NCX*_, *k*_*s*_ and *d* should all be unit order with respect to *N*. However, this cancellation is not possible for the calcium dynamics, as the NCX flux term is the only term that dependents on *N*. The kinetic constants *k, k*_*PMCA*_ and *g* must hence all be of comparable order as *N*. In the usual null-cline analysis, these rate constants are formally rescaled to unit order by *N*_*ext*_ and introduce a stiff parameter *γ* ≫ 1. We will instead insert *γ* directly and work with unit order kinetic parameters. This produces a fast calcium dynamic due to coupling with NCX, with coordinated 3:1 large sodium flux, and large ER calcium release rate.

The nullcline of the Calcium dynamics in the C-N phase plane produces an S-hysteresis with respect to the intracellular sodium concentration N, with a high-C branch and a low-C branch connected via a middle branch. The high-C turning point correspond to the NCX optimum calcium efflux and the low-C turning point corresponds to the ER optimum calcium release. The calcium dynamics undergo discontinuous jumps at these two turning points.

The nullcline of the Na dynamics produces a parabola in the C-N phase plane, with the maximum corresponding to the NCX influx of sodium. Oscillatory spiking occurs when this Na nullcline intersects the middle branch of the S hysteresis from the calcium null cline **(Figure 4F-G)**. The parameters were chosen at arbitrary units, at 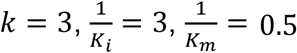, *k*_*PMCA*_ = 0.8, *C*_*K*_ = 0.6, *k*_*NCX*_ = 5, *K*_*a*_ = 0.35, *k*_*s*_ = 0.06, *d* = 0.2, *g* = 0.6, and [*Na*]_*ext*_ = 32. Initial conditions were set consistently at [*Na*]_0_ = 30 and [*Ca*]_0_ = 0.7. We note that when the ER optimum and the NCX optimum align, the parabolic null cline of Na is nearly tangent to the low-C branch of the Ca null cline hysteresis at the lower turning point. Hence, at this low-C branch, both the calcium dynamics and the sodium dynamics are slow, with the sodium concentration increasing gradually. In contrast, the two null clines are well-separated at the high-C branch, and the faster dynamics there correspond to the spike itself. Since tangency of the two null-cline curves corresponds to a very unique set of parameters, we expect the interval between spikes to be sensitive to change in the system parameters. In contrast, the spike itself on the high-C branch should be insensitive to stimuli or parameter changes. We see this in the model sensitivity analysis, where the peak heights are not sensitive to any parameter change, but oscillatory frequency is sensitive to change **(Figure 4H)**.

Interestingly, this calcium oscillation behavior is only possible when the IP_3_R and NCX-rate curves have similar peaks with the peak of IP_3_R-based calcium flux occurring at a slightly lower calcium concentration **(Figure 4I)**. For near tangency at the lower Calcium concentration branch with only one intersection at the middle branch of the Calcium nullcline, the larger bandwidth IP_3_R stipulates a larger [*Ca*^2+^]_*max*_ than NCX. Mathematically, this allows the small net Na and Ca influxes due to near-tangency to remain positive along the lower branch (the long low-calcium interval). This will eventually trigger the ER calcium spiking at the turning point since the calcium concentration will increase towards the [*Ca*^2+^]_*max*_ of the IP3 pathway. This ER-dormant interval hence relies on the bandwidths and amplitudes of the maxima of the IP_3_R and NCX pathways, which are related to the number and affinity of calcium binding sites on them.

### 2.6 Mathematical Model predicts cellular response to perturbations

We verified that this null-cline tangency model, due to alignment of calcium concentrations with high ER and NCX fluxes, can describe the experimental results we have discussed previously **(Figure 5A)**. It is known that VEGF primarily affects calcium signaling through IP_3_R. We demonstrated that the frequency of the calcium oscillation can be increased solely by modifying the 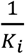 parameter from 3 to 3.3 **(Figures 5B-C)**. IP_3_ modulates IP_3_R-calcium release by inducing conformational changes that affect the allosteric calcium binding, which concurs with the model prediction that lower allosteric inhibitory calcium-IP_3_R binding will increase the spiking frequency.^67^

**Figure 5.**
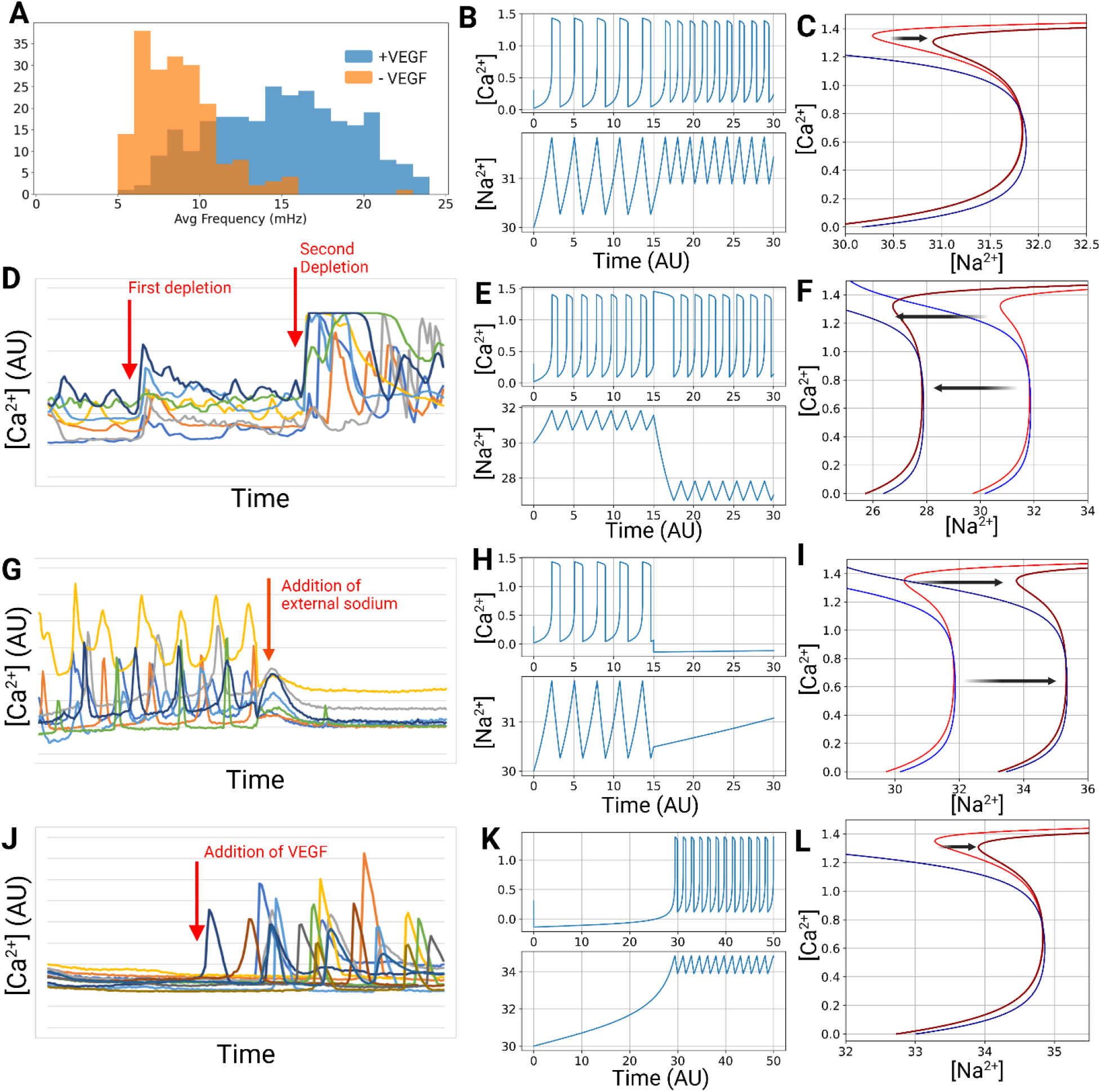
The NCX-IP_3_R model predicts the effects of external perturbations on calcium oscillation patterns. A. Overlaid histograms of the average oscillatory frequencies of ECs treated with and without VEGF, replicated from Figure 2. **B-C**. Oscillatory traces and phase plots with the parameter shift of 1/Ki from 3 to 3.3 introduced at timepoint 25. Calcium nullcline’s hysteresis peak shifts to the right, leading to a narrower oscillation zone and therefore higher frequency. **D**. Representative calcium concentration traces of several cells in a depletion setup which undergoes two rounds of depletion at the marked timepoints. During the depletion, cells exhibit a high calcium phase that is sustained for longer than a typical calcium oscillation peak. **E-F**. Oscillatory traces and phase plots with the parameter shift of [Na+]_ext_ from 32 to 28 introduced at timepoint 25. Both nullclines shift to the left, leading to a temporary phase in which cells are stuck in the high-calcium state during the transition phase, followed by resumption of oscillatory activity. **G**. Representative calcium concentration traces of several cells treated with sudden spike in external sodium concentration. Immediately following the addition of external sodium, all spiking activity ceases. **H-I**. Oscillatory traces and phase plots with the parameter shift of [Na^+^]_ext_ from 32 to 38 introduced at timepoint 25. Both nullclines shift to the right, leading to the peak of the calcium nullcline hysteresis extending beyond the bottom branch of the sodium hysteresis, which shifts the system to a non-oscillatory state. **J**. Representative calcium concentration traces of several cells which have been exposed to high external sodium concentration for several minutes before VEGF was introduced. The timepoint at which VEGF is added is marked with the red arrow. Sodium concentration remains at 300mM throughout the entire time. **K-L**. Oscillatory traces and phase plots with the [Na^+^]_ext_ set to 36 and 1/Ki parameter shifted from 3 to 3.3 at timepoint 35. The calcium hysteresis’s peak is shifted to the right, leading to the peak now extending no farther than the lower branch of the sodium nullcline, leading to resumption of oscillatory state. For figures C, F, I, and L, the dark red and dark blue lines represent the nullclines after the perturbation.

We also verified that the model exhibits the same calcium dynamics we observed with external sodium concentration modulation. First, when we adjust the external sodium concentration from 30 to 28, we see that the oscillation pattern is maintained, but crucially, there is a period of sustained high calcium concentration which is followed by continued oscillation **(Figure 5E-F)**. This reflects the behavior of endothelial cells during external cation depletion **(Figure 5D)**. Crucially, this behavior is observed even when the amplitude of change in external ion concentration is relatively small similar.

We also verified that the sudden spike in external sodium from 30 to 38 can lead to a sudden stop of oscillations and the cells being stuck in the “low calcium” state due to the lack of an oscillatory steady state point **(Figure 5H-I)**. This again matches the experimental data from external sodium spiking **(Figure 5G)**.

Lastly, we demonstrated that with the addition of external sodium, the two nullclines are shifted to the right which leads to a suspension of oscillation. Therefore, when we introduce to the system which already lacks oscillation due to high external sodium 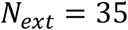 the 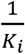 parameter change from 3 to 3.3 simulating VEGF addition, the model predicts a resumption of oscillation due to the calcium hysteresis shifting back further right leading to the peak of the hysteresis to shift further than the lower branch of the sodium nullcline (**Figure 5K-L**). We experimentally validate this by introducing 50ng/mL of VEGF into cell population with the external sodium spiked to 250 mM while keeping the sodium concentration constant. This results in the resumption of oscillatory activity, validating the model predictions (**Figure 5J, Supplementary Movie S13**).

## 3. Discussion

Regulation of intracellular calcium dynamics is a crucial signaling mechanism that can stimulate a diverse array of cellular behaviors in all eukaryotic cell types. Calcium signaling occurs in a wide range of frequencies, from the high-frequency oscillations in excitable cells such as muscle cells, or the gradual changes in cytosolic levels in various epithelial cells. Between these two ends lies non-excitable but calcium-oscillatory cells, most notably the endothelial cells, which undergoes cycles of rapid calcium release into the cytoplasm from the endoplasmic reticulum. These calcium oscillatory activities have been shown to play a crucial role in regulating endothelial behavior such as angiogenesis. However, these calcium oscillations in endothelial cells are more irregular and noisier compared to that of excitable cells, making it challenging to extract and analyze the data at a large population-wide scale. Because of this, it is yet unclear how the endothelial cells modulate the calcium signaling patterns when introduced to external stimuli.

To this end, we developed a computational pipeline that can extract features such as frequency and cross-correlation in endothelial cells in a robust manner. Using this computational approach, we uncovered unique transient and chronic responses of ECs to pro-angiogenic VEGF treatment, where the cells undergo a synchronization event within seconds of VEGF introduction, followed by a long-term shift in oscillation frequency. We have shown that this shift in frequency in response to VEGF-activation is replicated in iPSC-derived endothelial cells as they become mature and express sufficient levels of VEGF, indicating that this frequency shift due to VEGF signaling is highly conserved in endothelial lineage cells and highlights a potential signaling mechanism that regulates angiogenesis.

To take advantage of this mechanism, we designed a microfluidic channel to deplete the cations from the cell media to confirm that calcium spiking behavior can be manipulated through external ionic changes. We used external electrical current to induce the cell synchronization event similar to that caused by VEGF, which led to similar transcriptional changes in genes downstream of VEGF that are involved in angiogenesis. These results suggest that ECs respond to VEGF signal with a unique calcium signaling pattern which can be mimicked using electrical currents, which can potentially be used for precise spatiotemporal control of angiogenic processes that can replace or augment traditional growth factor treatments.

Inspired by these results, we hypothesized that calcium oscillation is coupled with sodium ions in the non-excitable ECs. We developed a mathematical model based on mechanisms of NCX transporter and IP_3_R-based calcium release from the endoplasmic reticulum and found that our model creates oscillatory calcium patterns that we observed in endothelial cells. We then confirmed that the model can accurately predict the EC responses to various external stimuli such as VEGF stimulation, external ion depletion or supplementation, demonstrating the potential value of our mathematical model for explaining calcium behavior in endothelial cells.

Numerous mathematical models have been proposed to describe the calcium spiking behavior in various excitable cells such as the Hodgkin-Huxley model or the Fitzhugh-Nagumo model. However, these existing models focus on the coupling of the membrane potential and calcium concentration to explain the rapid oscillation seen in excitable cells, making them inadequate for modeling the calcium oscillation behavior in non-excitable cells where the rate of change of membrane potential is significantly slower than that of calcium. Therefore, we propose here a novel mathematical model that can replicate the oscillatory system without relying on the coupling of membrane potential with calcium, instead relying on the similar activation-inhibition curve of NCX-transporter and IP3R with respect to intracellular calcium concentration. We show here that our model faithfully models the rapid calcium changes without a similarly large magnitude of fluctuations in sodium, which explains why membrane potential does not fluctuate at a significant magnitude. We also show that when we introduce perturbations to the parameters in the model that correspond to experimental stimulation (i.e. VEGF, depletion, external sodium), our model accurately predicts the cellular behavior. Therefore, our model can potentially be used to predict how cells may react to calcium-affecting drugs, electrical stimulations, and others.

In summary, we report that endothelial cells modulate the calcium spiking behavior via frequency and cross-correlation in response to angiogenic signals. We have shown that we can mimic the effects of VEGF via direct manipulation of calcium signaling through carefully guided electric currents within our microfluidic setup. We mathematically capture these behaviors with a novel concept of near-tangent null clines that reflects the cross-talk between activated/inhibited NCX and IP3R pathways. Through the predictive mathematical model and electric stimulation proposed in this study, future studies should explore manipulation of angiogenic cationic responses independent of needing VEGF delivery and 3D tube formation in angiogenic sprouting assays in controlled cation environments. These findings suggest that precise control of endothelial calcium dynamics may serve as a powerful strategy for engineering vascular behaviors in tissue models and regenerative systems.

## 4. Methods

### Human iPSC and BEC Culture

Human BECs derived from the dermis of an adult donor (Promocell) were expanded and used between passages 5 and 8. The cells were cultured in MV2 media (Promocell) and similarly incubated at 37°C with 5% CO_2_. Human iPSCs derived from lung fibroblast (WiCell) were seeded on tissue culture plastic surfaces coated with 0.083 mg/mL of Matrigel (Corning) in DMEM-F12 media (Gibco) and grown in mTeSR+ media (Stem Cell Technologies) at 37°C with 5% CO_2_. 10 μM of Y-27632 Rho-kinase inhibitor was added to the cell culture media for the first day after seeding, then subsequently removed. All cell lines were routinely tested for mycoplasma contamination and were negative throughout this study.

### Human iPSC Differentiation to BEC

Human iPSCs were seeded in single cell suspension at 30,000 cells/cm^2^ in mTeSR+ media containing 10 μM of Y-27632 for one day. The media was switched to 4mL of APEL2 media (Stem Cell Technologies) containing 6 μM of CHIR99021 (Stem Cell Technologies) per well in a 6-well plate, and allowed to differentiate for 2 days. Following this, the media was switched to 4mL of APEL2 media containing 50ng/mL of VEGF-A (Stem Cell Technologies), 25ng/mL BMP4 (Stem Cell Technologies), and 10 ng/mL of FGF2 (R&D Biosciences) and left for two days. Following this, the cells were detached using TrypLE Express (Gibco) and replated at a density of 50,000 cells/cm^2^. The cells were cultured in MV2 media containing 50ng/mL of VEGF-A for up to 4 days with media change every 2 days.

### Calbryte 520AM Staining and Imaging

Calbryte 520AM dye (AAT Bioquest) resuspended in DMSO was added to serum-free MV2 media at a final concentration of 5 μM. The cells were washed 1x with dPBS without calcium, then incubated with the Calbryte staining solution for 60 minutes. Following the incubation, the staining solution was removed and the cells were washed 1x with dPBS without calcium. Fresh media was added, then the cells were imaged at interval of 5 seconds with GFP excitation laser. For fluorescence imaging, either the Agilent Lionheart microscope (BioTek) or the A1R confocal microscope (Nikon) were used at 10x objectives.

### Electrical Stimulation

Human BECs were seeded in either Ibidi single channel microfluidic chip (Ibidi) or a 6-well plate and allowed to reach 100% confluency. SourceMeter 2400 source measure unit (Keithley) was used to generate the electrical pulse current. The electrodes were placed on either end of the channel for the microfluidic chip, or on the opposite edges of the well in a 6 well plate. Voltage values were held constant between 10-12 V and the current reached 10-12 mA across the Ibidi chip. Since the well plate experiences different resistance depending on the volume of cell culture media, we empirically tested various media volumes until we determined the correct volume to reach the same resistance across the well as the Ibidi chip, at 950μL of MV2 media. The chip or the well plate connected to the electrode was placed inside Agilent Lionheart microscope’s incubation chamber and the imaging was started at least 5 minutes prior to applying the electrical stimulation to measure the transient effects.

### Gene Expression

To analyze the gene expressions, we collected the RNA from BECs using Qiagen Mini-prep kit (Qiagen), and quantified and diluted to the same concentration for all samples. The RNA was reverse transcribed using High-capacity cDNA reverse transcription kit (Thermo Fisher) using the recommended protocol from the manufacturer. We used TaqMan Gene expression assays (Thermo Fisher) following manufacturer protocols and used GAPDH as endogenous controls to normalize relative gene expression levels using the ΔΔC_T_ method. All statistical analysis was done on the ΔΔC_T_ values.

### Nondimensionalization of the model

Consider the dimensional equation for calcium and sodium below.

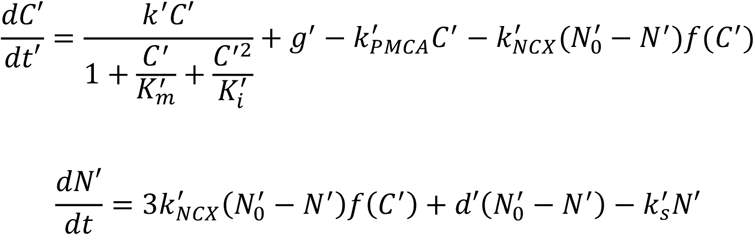

We note *k*_*s*_ has units of inverse time. Since all the fluxes for sodium should be the same order and unit order with respect to the Calcium fluxes, we can choose *1/k*_*s\**_ as the characteristic time. We define 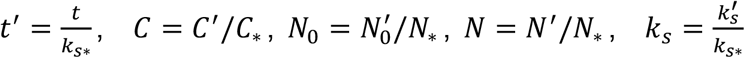, where the unprimed quantities are dimensionless. Typical values we observe would be *C*_∗_ = 1 *μM*, and *N*_∗_ = 5 − 10 *mM*, making 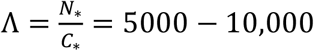. Then, the sodium equation becomes

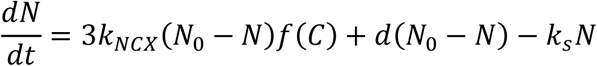

where 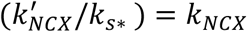 and 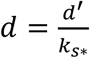 are all unit order dimensionless parameters.

The calcium gating function for NCX is 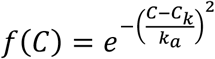 where we have used 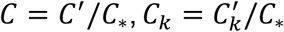, and 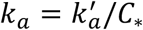. Using the same scaling, we get

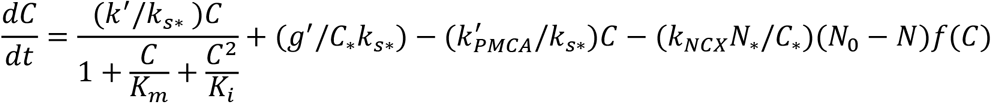

We note that the large If we define Λ = *N*_∗_/*C*_∗_ term that appears in the NCX flux term for calcium that does not appear in the sodium N equation. To balance this term, all the other dimensionless flux kinetic coefficients 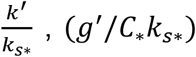 and 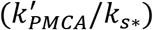 should also be large of order Λ. We hence rescale them to extract the lone large parameter Λ and render all the parameters of order unity with respect to Λ:

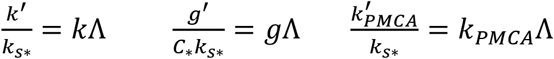

where *k, g* and *k*_*PMCA*_ are now all unity order.

We get the dimensionless equation with a stiff dimensionless parameter and with dimensionless variables, time and parameters.

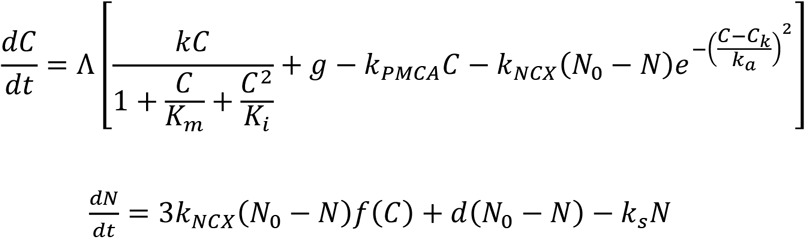

where the stiff parameter is Λ = *N*_∗_/*C*_∗_, then all the kinetic parameters *k, g, k*_*PMCA*_ and *k*_*NCX*_ are dimensionless and of unit order with respect to Λ.

Note, with this order assignment, we assume that *k’ >> k*_*s*_, *g*^′^ ≫ *C*_∗_*k*_*s*_ and 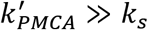 but 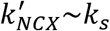, which is reasonable considering the high Calcium concentration spikes require large calcium fluxes, much larger than sodium fluxes. Also note that because of the difference in equilibrium calcium and sodium concentrations, the NCX flux produces large fraction change in the calcium concentrations but not the sodium concentrations. This is the key to stiff dynamics with large Λ to allow the quasi-steady state nullcline analysis.

## Acknowledgements

We acknowledge support from the University of Notre Dame through “Advancing Our Vision” Initiative in Stem Cell Research, Harper Cancer Research Institute – American Cancer Society Institutional Research Grant (IRG-17-182-04), American Heart Association through Career Development Award (19-CDA-34630012 to D.H.-P.), National Science Foundation (2047903 and 2225601 to D.H-P., 2120200 to EMBRIO Institute), National Institutes of Health (R35-GM-143055 to D.H.-P. and R35-GM-156615 to J.Z.), National Science Foundation-Graduate Fellowships Program (NSF-GRFP to D.P.J.).

## Data and Code Availability

The relevant imaging data are present in the Supplementary Information and Supplementary Movies. Raw images are deposited in FigShare. The original code used in this study can be found on Github Repository [https://github.com/donghyunjeong21/calcium_signaling].

