## Supplementary Materials for "Harnessing NCX-IP_3_R-dependent Calcium Oscillations to Regulate Angiogenic Signaling in Endothelial Cells"

**This PDF file includes:**

Supplementary Figures S1 to S4

**Other Supplementary Materials for this manuscript include the following:**

Movies S1 to S13

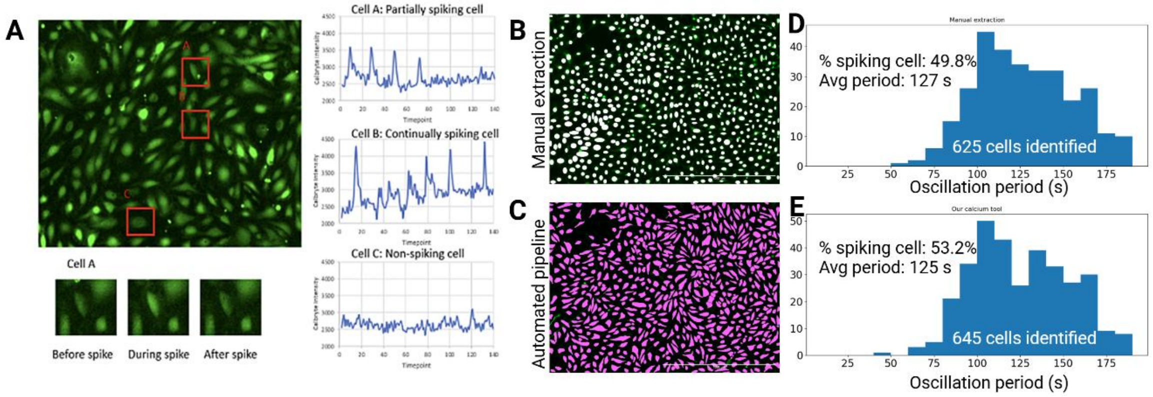


**Supplemental Figure S1.** Comparison of the automated calcium data analysis tool with manual segmentation and extraction. **A.** An example of a calcium staining image with 3 sample cells identified as partially, continuously, and non-spiking cells. **B-C.** Cell segmentation results from manual ROI identification and Cellpose-based automated algorithm. **D-E.** Distribution of average oscillation period in the cells identified by each method.

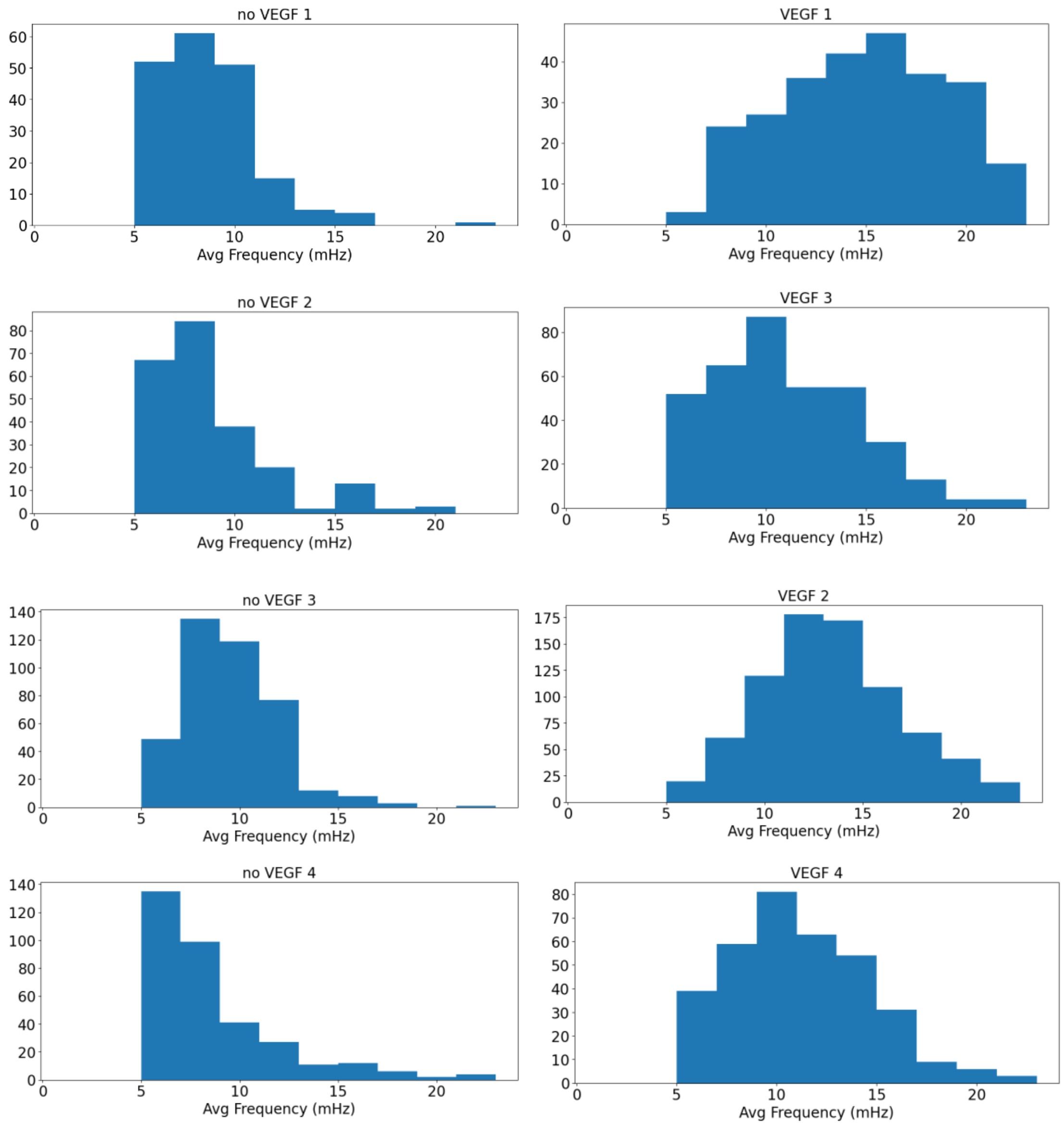


**Supplemental Figure S2.** Calcium oscillation frequency shift when treated with VEGF. Each histogram consists of cells collected from a distinct experimental setup with cells cultured on separate days. Each VEGF-no VEGF pair was run on the same day.

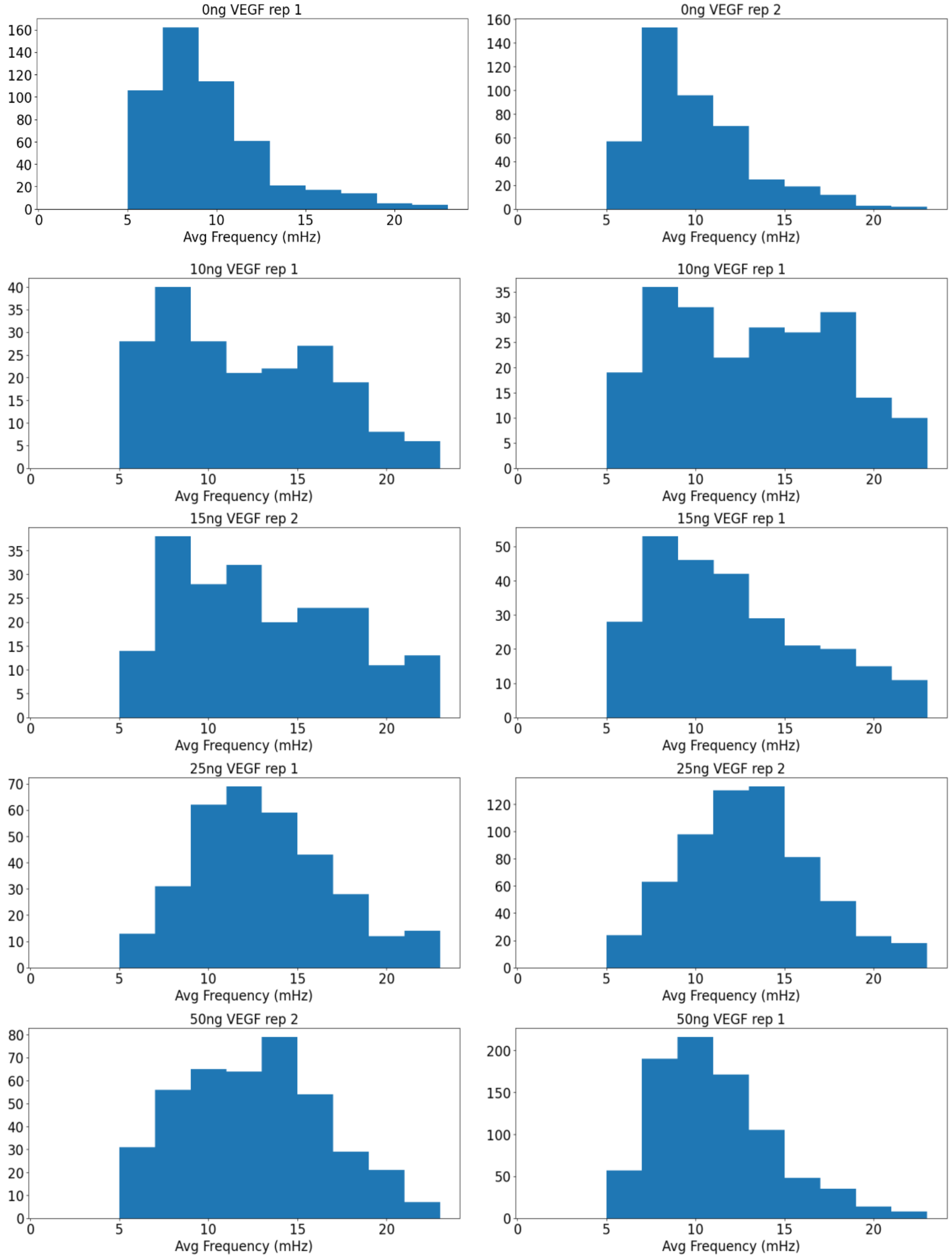


**Supplemental Figure S3.** Calcium oscillation frequency shift when treated with various concentrations of VEGF. Each concentration was tested in duplicates (n=2). All concentrations tested above 10ng/mL resulted in the oscillation frequency shift.

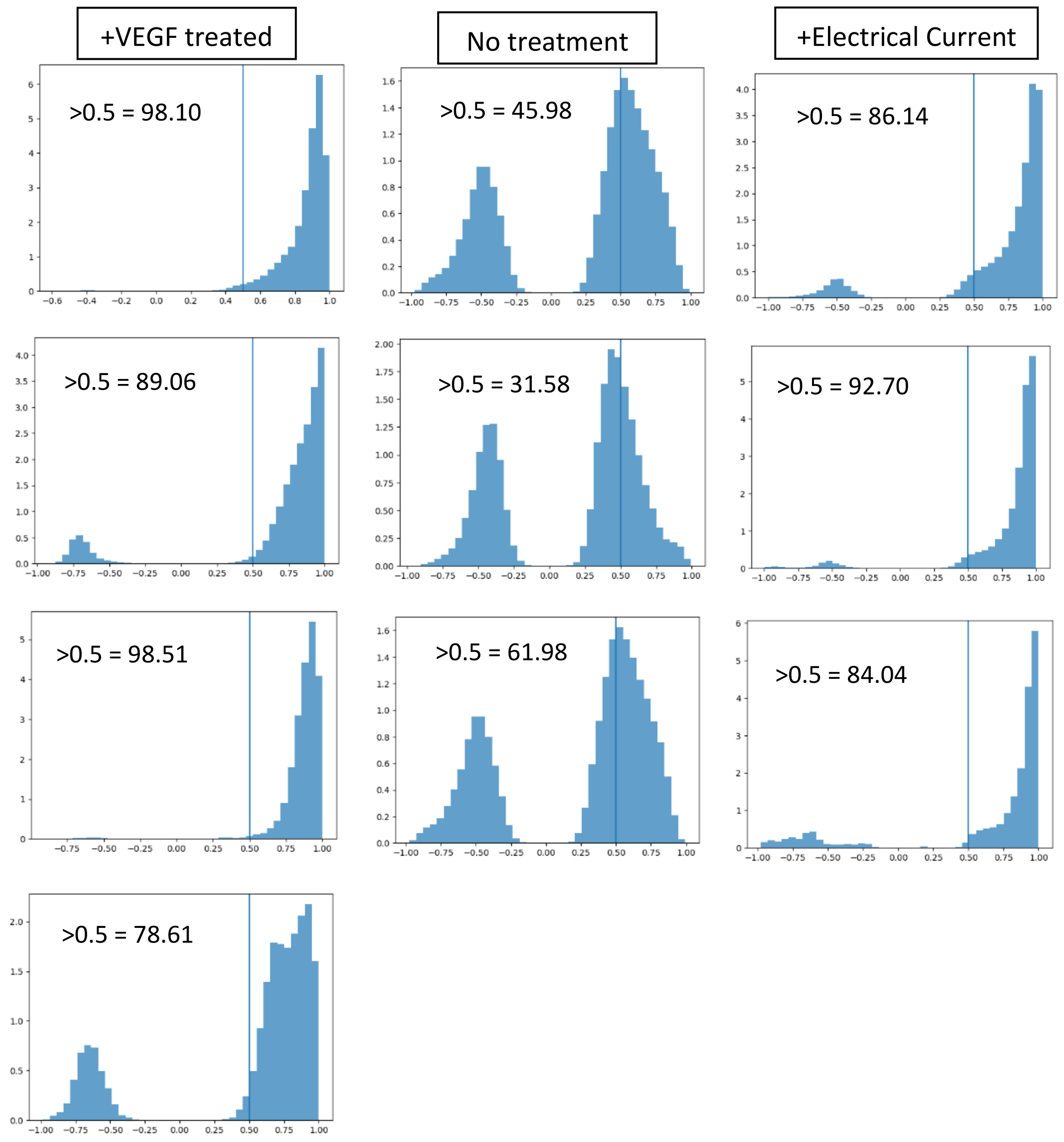


**Supplemental Figure S4.** Histograms of cross-correlation values for each cell-cell pair within the imaging window. Left column shows cell correlation immediately following treatment with 50ng/mL of VEGF. Right column shows cell correlation during electric current treatment of 10mA and 5V. The center column shows correlation for untreated cells undergoing natural oscillation. N=4 for VEGF and n=3 for control and electric current.
